# A novel method for fine-scale retrospective isotope analysis in mammals using eye lenses

**DOI:** 10.1101/2024.02.29.582866

**Authors:** Kazuki Miura, Jun Matsubayashi, Chikage Yoshimizu, Hino Takafumi, Yuri Shirane, Tsutomu Mano, Hifumi Tsuruga, Ichiro Tayasu

## Abstract

1. Investigating individual behavioural variations in mammals is essential for understanding their ecology and evolution, and plays a critical role in conservation and management practices. However, the reconstruction of long-term individual behaviour, such as via bio-logging, remains challenging owing to cost constraints and limitations of battery life and the impact of device size for smaller animals. This study proposed and tested a novel and cost-effective method for retrospective isotope analysis using mammalian eye lenses, specifically focusing on brown bears (*Ursus arctos*).

2. Seven pairs of bear eye lenses were collected from southwestern Hokkaido, Japan. One or both lenses of each bear were segregated into small fragments from the outermost to the core tissues, and the nitrogen and carbon stable isotope ratios (δ^15^N and δ^13^C) of each eye lens fragment were measured. These isotope ratios in the eye lenses were compared with the δ^15^N and δ^13^C of potential brown bear diets in the study region (C_3_ herbs, C_3_ fruits, terrestrial animals, and corn). Additionally, we compared the isotopic patterns in both the right and left lenses of the same bear in two individuals to evaluate the consistency of our preparation protocol.

3. In all eye lenses, high δ^15^N values were identified near the core, which gradually decreased towards the outer tissues, indicating ontogenetic dietary shifts related to lactation and weaning in the early life stage. Bears from study areas with high corn availability exhibited substantial increases in δ^13^C and δ^15^N in the outer lens tissues, suggesting a dietary shift toward corn consumption after weaning. Isotopic patterns between lens pairs from the same individual were similar, although discrepancies increased in tissues located 1.00 to 2.25 mm from the core, highlighting the need for standardisation in sample processing.

4. This study demonstrates a novel and simple technique for retrospective isotope analysis in wild mammals using eye lenses, effectively reconstructing the feeding histories of brown bears. Our findings provide a new avenue for studying individual time-series behavioural patterns, with important implications for the fields of mammalian ecology, evolution, conservation, and management.

## 1. Introduction

Knowledge of behaviour is fundamental to understanding animal ecology and evolution. Most previous animal behaviour studies have targeted interspecific and population-level behavioural variations (Wolf & Weissing, 2012). In contrast, recent studies have highlighted the importance of understanding individual behavioural variations, which play a crucial role in ecological adaptations, inter- and intra-specific interactions, and population and community dynamics (Araújo et al., 2011; Newsome et al., 2015; Wolf & Weissing, 2012). This information also has profound implications for the conservation and management of wild animals (Honda et al., 2018; Swan et al., 2017).

Mammals demonstrate marked individual behavioural variations (Dickman & Newsome, 2015; Swan et al., 2017). Historically, the identification of individual behavioural variations in mammals has relied mainly on direct observation (Shimozuru et al., 2017), DNA analysis (via repeatedly capturing individuals or collecting animal tissues; Berezowska-Cnota et al., 2023), and bio-logging using electrical devices such as radio, GPS, accelerometers, and animal-borne sensors (Hertel et al., 2021; Naganuma et al., 2021). However, these methods have several limitations. For instance, direct observation is difficult for many mammals because they are often human-averse, highly mobile, and inhabit environments that are not easily visible (Cagnacci et al., 2010; Newsome et al., 2009). Repeatedly capturing individuals or collecting animal tissues for DNA analysis over a wide spatial area and extended periods involves high human and financial costs (Shimozuru et al., 2022). Bio-logging techniques potentially overcome these challenges, but are not easily applicable for a large number of individuals and/or a wide array of mammalian species, given the body-size limitations for equipped mammals, high financial cost, and substantial efforts required for non-lethal capturing and recapturing of individuals (Cagnacci et al., 2010).

To address these challenges, recent studies have increasingly emphasised the effectiveness of retrospective isotope analysis in examining individual-level behaviours, such as feeding and movement patterns (i.e. “iso-logging”; Matsubayashi et al., 2022). This approach employs incremental and metabolically inert tissues, such as the hair, claws, and teeth of mammals, to reconstruct temporal isotopic changes within the same individual (Cerling et al., 2006; Newsome et al., 2009, 2015). Although these methods have contributed to our understanding of individual behavioural patterns (Cerling et al., 2006; Hata et al., 2017; Newsome et al., 2015), there are notable limitations. First, traditional methods using hair and claw specimens for reconstructing isotopic histories are limited to time-series of a few months or years due to tissue shedding (Ethier et al., 2010; Jimbo et al., 2020). Second, the tooth dentin method, which is capable of reconstructing lifetime multi-element isotopic ratios (Hobson & Sease, 1998; Zazzo et al., 2006), often requires specialised pre-treatment and detailed knowledge on the growth pattern for each type of tooth, restricting its widespread use (Czermak et al., 2020; Waugh et al., 2018). Therefore, simpler methods to reconstruct lifetime changes in isotope ratios without special techniques would substantially advance the field of mammalian individual behavioural research.

Recent isotopic studies have demonstrated the potential of animal eye lenses for retrospective isotope analyses (Bell-Tilcock et al., 2021; Matsubayashi et al., 2022; Wallace et al., 2014). Eye lenses, primarily composed of structural proteins called crystallins, grow incrementally, and new tissues are sequentially added to the outermost part throughout an individual’s lifetime (Peebles & Hollander, 2020; Wride, 2011). Once these tissues are formed, protein synthesis ceases, and they no longer experience metabolic turnover (Peebles & Hollander, 2020). Thus, longitudinal tissue segregation and isotope analysis of eye lenses can enable the reconstruction of individual behavioural histories from isotopic chronology (Bell-Tilcock et al., 2021; Matsubayashi et al., 2022; Peebles & Hollander, 2020). This method has successfully been applied in fish and cephalopods (Hunsicker et al., 2010; Wallace et al., 2014); however, its applicability and appropriate lens segregation methods in mammals have not yet been tested.

In this study, we established a novel experimental protocol for longitudinally segregating eye lenses from wild mammals and investigated whether retrospective isotope analysis of eye lenses could reconstruct ontogenetic dietary shifts. Individual diet history is an important indicator of feeding behaviour. We focused on the nitrogen and carbon stable isotope ratios (δ^15^N and δ^13^C, respectively) in the eye lenses of brown bears (*Ursus arctos*) in Hokkaido, Japan. Omnivorous brown bears exhibit larger individual dietary variations than other mammals, such as humans (Kusaka et al., 2016; Matsubayashi et al., 2015), and live for over 20 years (Hobson et al., 2000). δ^15^N is a valuable indicator of dietary shifts in mammals, as they theoretically reflect enriched δ^15^N in tissues during lactation (Hobson et al., 2000; Newsome et al., 2009). We hypothesised that bear eye lenses retain isotopic information from the infantile period and predicted that a gradual decrease in δ^15^N would be observed from the core of the lens toward the outer tissues, reflecting the weaning process.

We also assessed the potential of δ^13^C in bear eye lenses to track dietary shifts over time. δ^13^C is a well-established indicator of C_4_ plant consumption, including corn (*Zea mays*), and C_4_ plant-based diets lead to significantly higher δ^13^C than other bear diets (Ditmer et al., 2016; Matsubayashi et al., 2015). We predicted that corn consumption would be reflected by a substantial increase in δ^13^C within specific sections of the eye lens, assuming the preservation of isotopic ratios. Notably, corn (mainly dent corn) is the only C_4_ plant available to brown bears in Hokkaido, Japan, and its consumption primarily occurs in late summer (Hata et al., 2017; Sato et al., 2005). Therefore, analysing δ^13^C changes in eye lens sections could provide valuable insights into both the preservability and temporal resolution of the isotopic information recorded in mammalian eye lenses.

This study investigated three key questions via retrospective nitrogen and carbon stable isotope analyses of eye lens samples from Hokkaido brown bears. First, we explored whether δ^15^N of the eye lens nucleus could reconstruct lactation and weaning patterns. Second, we examined whether the δ^13^C of each eye lens section could reveal the onset and seasonal variation of corn-feeding signatures. Finally, by comparing the isotopic patterns of left and right eye lenses from the same individual, we assessed whether our preprocessing method could reliably reproduce similar isotopic patterns in both eyes (i.e. assess the reproducibility of the isotope ratio chronology by our preprocessing protocol). Via these examinations, we aimed to elucidate the effectiveness of continuous eye lens isotope analysis in reconstructing the individual behavioural history of mammals.

## 2. Materials and Methods

### 2.1 Study site and sample collection

The study used bears from three towns (Assabu, Kuromatsunai, and Yakumo) located in or at the edge of the Oshima Peninsula, southwestern Hokkaido, Japan (Fig. 1). The Oshima Peninsula had an estimated bear population of 2,040 in 2021 (Mano et al., in press). This population is genetically distinct from those in the central and western Hokkaido regions (Matsuhashi et al., 1999). The annual mean temperatures and precipitation were 10.3, 7.5, and 8.1℃ and 1230.7, 1485.0, and 1294.1 mm in Assabu (represented by the nearest weather observatory, Esashi), Kuromatsunai, and Yakumo, respectively (Japan Meteorological Agency, 2023). The primary diet of brown bears in this region comprises C_3_ plants (herbaceous plants, berries, and acorns), terrestrial animals (insects and deer), and crops (mainly dent corn); however, the contribution of deer is minimal among terrestrial animals (Sakiyama et al., 2021; Sato et al., 2005). Additionally, their consumption of marine-derived resources, such as salmon and marine mammals, is negligible (Matsubayashi et al., 2015; Sato et al., 2005). A previous study found that the bears inhabiting in this region include individuals that are highly dependent on corn (Sakiyama et al., 2021). This information is crucial for testing our second prediction concerning δ^13^C variations in bear eye lenses (see Introduction).

**Fig. 1.**
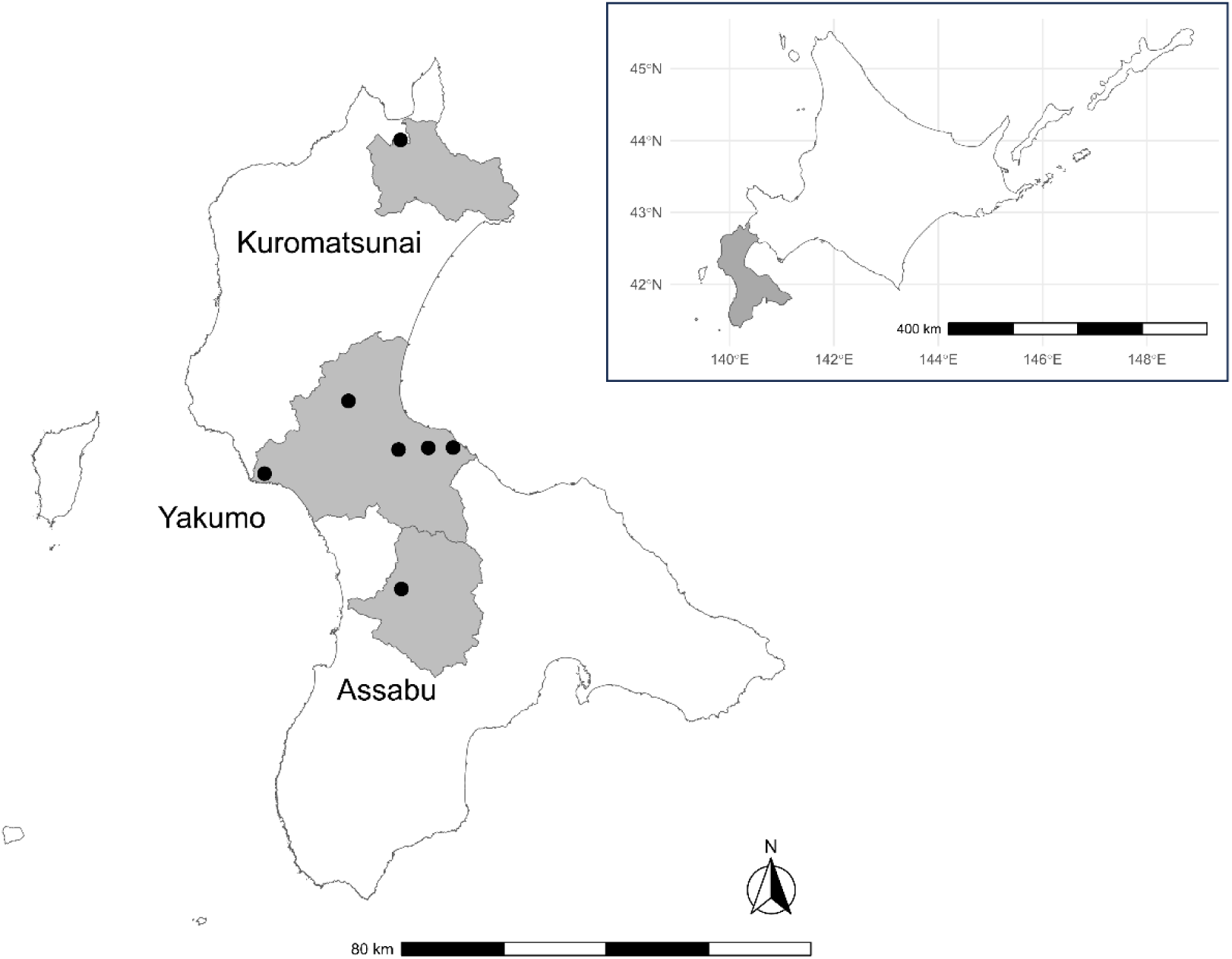
Geographical locations of the Oshima Peninsula, southwestern Hokkaido, Japan; the three towns; and harvest locations (black dots) of the seven included bears.

In the towns, the agricultural land cover ratios were 5.8, 12.2, and 7.8% in Assabu, Kuromatsunai, and Yakumo, respectively. In Yakumo specifically, dent corn is widely cultivated every year, and the total cornfield area and cover ratios of cornfields against total agricultural land were 781.3 ha and 10.4% in 2022 (unpublished data from the Yakumo Town Office), respectively. Few electrical fences have been constructed around the cornfields. Bear-related damage to the cornfields was observed annually (Fig. S1). In 2022, the damaged cornfield area totalled 10.1 ha, composing 1.3% of the total dent cornfield area (unpublished data from Yakumo Town Office). Given such high corn availability, samples of bears consuming substantial amounts of corn were expected from this town. In the other two towns, there was no prior information available on recent cornfield cover areas and the status of corn damage by bears. However, bear-related damage to cornfields has been observed in both towns over the past five years (Hokkaido Prefecture, 2024).

The Research Institute of Energy, Environment and Geology has collected biological samples of Hokkaido brown bears for monitoring purposes. Eyeballs were contained in some bear samples. Seven pairs of eyeballs were obtained from the above three towns: one from Assabu, one from Kuromatsunai, and five from Yakumo. All samples were harvested in 2022. Sex, location of death, and body length (cm from the tip of the nose to the anus) data were recorded for each individual (Table 1). The ages of the bears were determined by counting the cementum annuli on a fourth premolar (Yoneda, 1976). The collected eyeballs were preserved in a freezer at -25 °C until dissection.

**Table 1.**
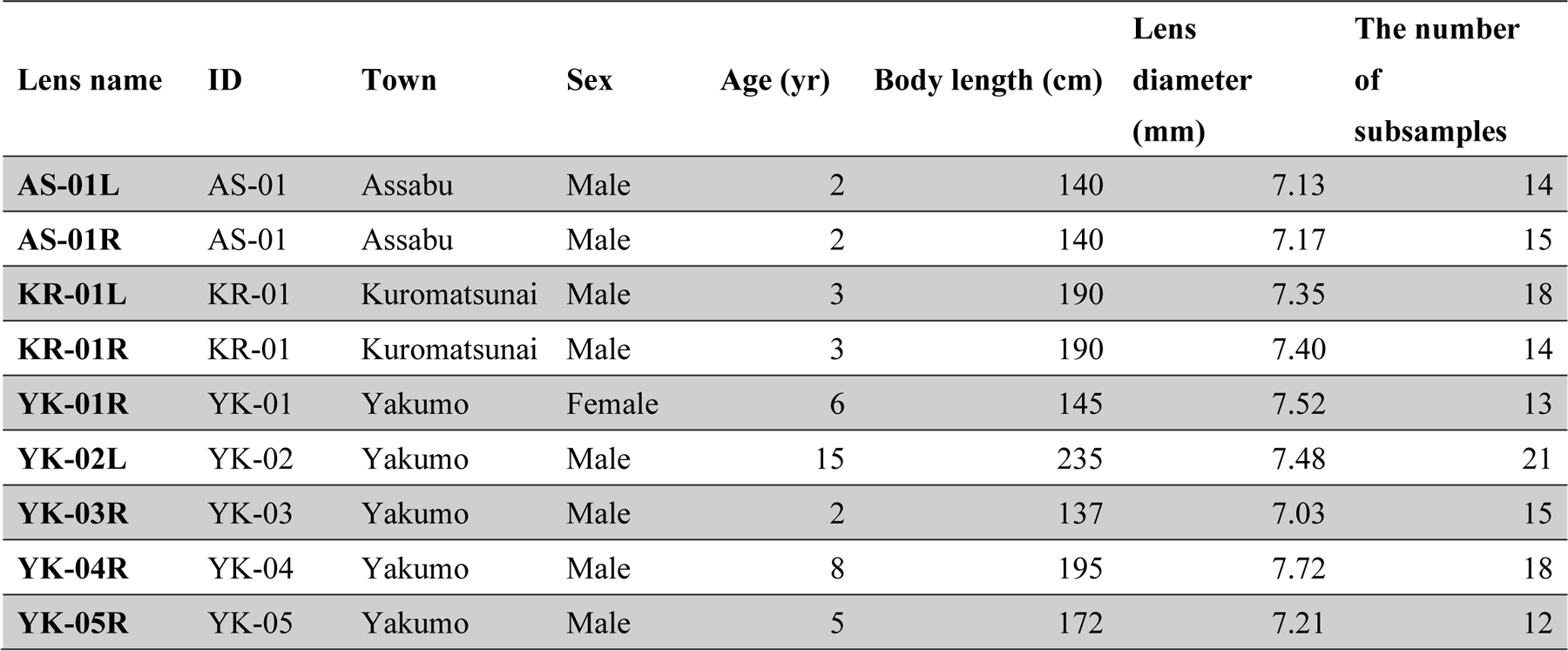
Summary of the information on targeted bear individuals, eye lenses, analysed subsamples of eye lens in the present study.

The biological samples used in the study were obtained from bears killed for nuisance control measures and game hunting in accordance with the governmental plan of Hokkaido Prefecture (Hokkaido Prefecture, 2022). Therefore, this study did not require specific ethical approval.

### 2.2 Eye lens preparation

For stable isotope analysis, this study used lenses of the right eye for most individuals (*n* = 7). Owing to prior allocation for the preliminary test, the left lens from the Yakumo sample (YK-02) was used instead. Additionally, left lenses from the Assabu and Kuromatsunai individuals (AS-01 and KR-01) were used to compare isotopic patterns within the same individual (*n* = 2). This resulted in a total of nine analysed eye lenses.

We developed a novel pretreatment protocol for drying and hardening the lenses before dividing them for sequential isotope analysis. Existing methods for dividing raw fish and cephalopod eye lenses (Hunsicker et al., 2010; Wallace et al., 2014) were not suitable for bear lenses because of their marked pliability and challenges in separation.

Our novel protocol comprised the following steps:

1. Eye lenses were meticulously dissected from the eyeballs using surgical scissors and tweezers, while retaining the covers of the vitreous humor and lens capsule, both of which are transparent gel-like substances mainly comprising water and collagen proteins (Danysh & Duncan, 2009; Mishra et al., 2023). If black suspensory ligaments and ciliary bodies were attached to these substances (Feher, 2012), they were also retained. This meticulous approach prevents potential lens damage arising from the forceful removal of these structures (Fig. S2).
2. Each dissected lens, still encapsulated by the vitreous humor and lens capsule, was positioned on a tungsten mesh net (1.5 × 1.5 mm mesh size) within a glass dish, ensuring no contact with the dish bottom.
3. As brown bear lenses possess a biconvex shape (DOĞAN et al., 2020), all lenses were oriented with the more prominent convexity facing upward. Drying then proceeded for at least 24 h at 35°C.

### 2.3 Eye lens segregation

After the eye lenses were dry, the vitreous humor and lens capsule, which covered the outermost portion of each dried lens, were meticulously removed using a pair of tweezers under a stereomicroscope. This was achieved via gradual maceration with distilled water. After removal of the vitreous humor and lens capsule, the long axis of the eye lens (i.e. the equatorial diameter) was measured with an electronic calliper to an accuracy of 0.01 mm, and the hemispherical portion of the lens was grasped and fixed with flat-tipped forceps. Eye lens segregation was performed as follows:

1. The entire surface of the lens hemisphere, opposite the hemisphere fixed with flat-tip forceps, was gradually shaved as evenly as possible using tweezers. This was achieved via gradual maceration with distilled water.
2. The shaved tissues were placed in one well of a 96-well plate (hereafter, “subsample”).
3. The equatorial diameter of the shaved eye lens was measured.
4. For a single-lens sample, steps 1–3 were repeated until the lens diameter was reduced to half of the initial measurement, reaching the radius of the equatorial diameter.
5. After lens segregation, the subsamples were redried at 35°C for at least 24 h.

### 2.4 Stable isotope analysis

Approximately 0.50 and 0.15 mg of the dried subsamples were placed in tin capsules for normal and sensitive analyses, respectively. The nitrogen and carbon stable isotope ratios were determined using a Delta V Advantage isotope ratio mass spectrometer (Thermo Fisher Scientific, Waltham, MA, USA) coupled with a Flash EA 1112 analyser (Thermo Fisher Scientific) via a Conflo IV interface (Thermo Fisher Scientific). The isotope ratios were denoted by δ values relative to an international standard scale as follows:

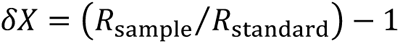

where *X* represents either ^15^N or ^13^C. *R*_sample_ is the ^13^C/^12^C or ^15^N/^14^N ratio of the sample, and *R*_standard_ refers to the ^13^C/^12^C ratio of Vienna Pee Dee Belemnite (VPDB) or ^15^N/^14^N ratio of atmospheric nitrogen. Isotope ratios were then corrected via two-point calibration using two laboratory standards (CERKU-02: δ^15^N = 22.71‰ and δ^13^C = -19.04‰, CERKU-03: δ^15^N = 2.18‰ and δ^13^C = -34.92‰) which are traceable to multiple international standards (Tayasu et al., 2011). The estimated analytical errors of δ^15^N and δ^13^C were within 0.11‰ and 0.06‰ for CERKU-02, and within 0.06‰ and 0.09 ‰ for CERKU-03, respectively. The lens subsamples were not defatted, as the lipid content in mammalian eye lenses is generally less than 1% (Feher, 2012).

The patterns of δ^15^N and δ^13^C, along with the distance from the core (mm), were illustrated in line graphs, with each individual depicted separately. The distance of each subsample from the core was represented as the midpoint of the shaved thickness of each subsample. Stable isotope ratios for potential dietary components of brown bears in the study region were obtained from Matsubayashi et al. (2015) and Sakiyama et al. (2021), and were shown in the graph. All illustrations of the isotopic patterns were performed using R 4.3.2 (R Core Team, 2023), with the ggplot2 ver. 3.4.4 package (Wickham, 2016).

## 3. Results

The body lengths of the brown bears ranged from 137 to 235 (mean ± SD: 173.43 ± 36.02) cm, and their ages were 2 to 15 (5.86 ± 4.60) years. One individual killed in Yakumo was a female, and all others were males (Table 1). The mean (±SD) equatorial diameter of the nine dried eye lenses was 7.33 (± 0.22) mm. The number of segregated subsamples ranged between 12 and 21 (15.56 ± 2.88). The mean segregated thickness between each subsample was 0.24 (± 0.13) mm. Two subsamples were not analysed because of the low sample volume (<0.15 mg). The carbon-to-nitrogen (C:N) atomic ratios of the analysed subsamples ranged between 3.56 and 3.86, with a mean of 3.66 (± 0.07).

The δ^15^N fluctuated significantly from the core to the outer subsamples in all seven lenses (Fig. 2a). The mean δ^15^N of the most central subsample that could be analysed in each lens was 5.4‰ (± 0.9; range between 4.1 and 6.5‰), which were segregated by approximately < 0.50 mm from the core in all seven lenses (Fig. 2a). All individuals showed a gradual decrease in δ^15^N from the core to approximately 2.50 mm from the core. The average minimum δ^15^N of each individual was 3.5‰ (± 1.0; range between 2.3 and 5.1‰), observed between the core and the subsample from 2.50 mm from the core. The average difference in δ^15^N between the central subsample and minimum value within a single eye lens was 1.9‰ (± 1.0; range from 0.5 to 3.3‰).

**Fig. 2.**
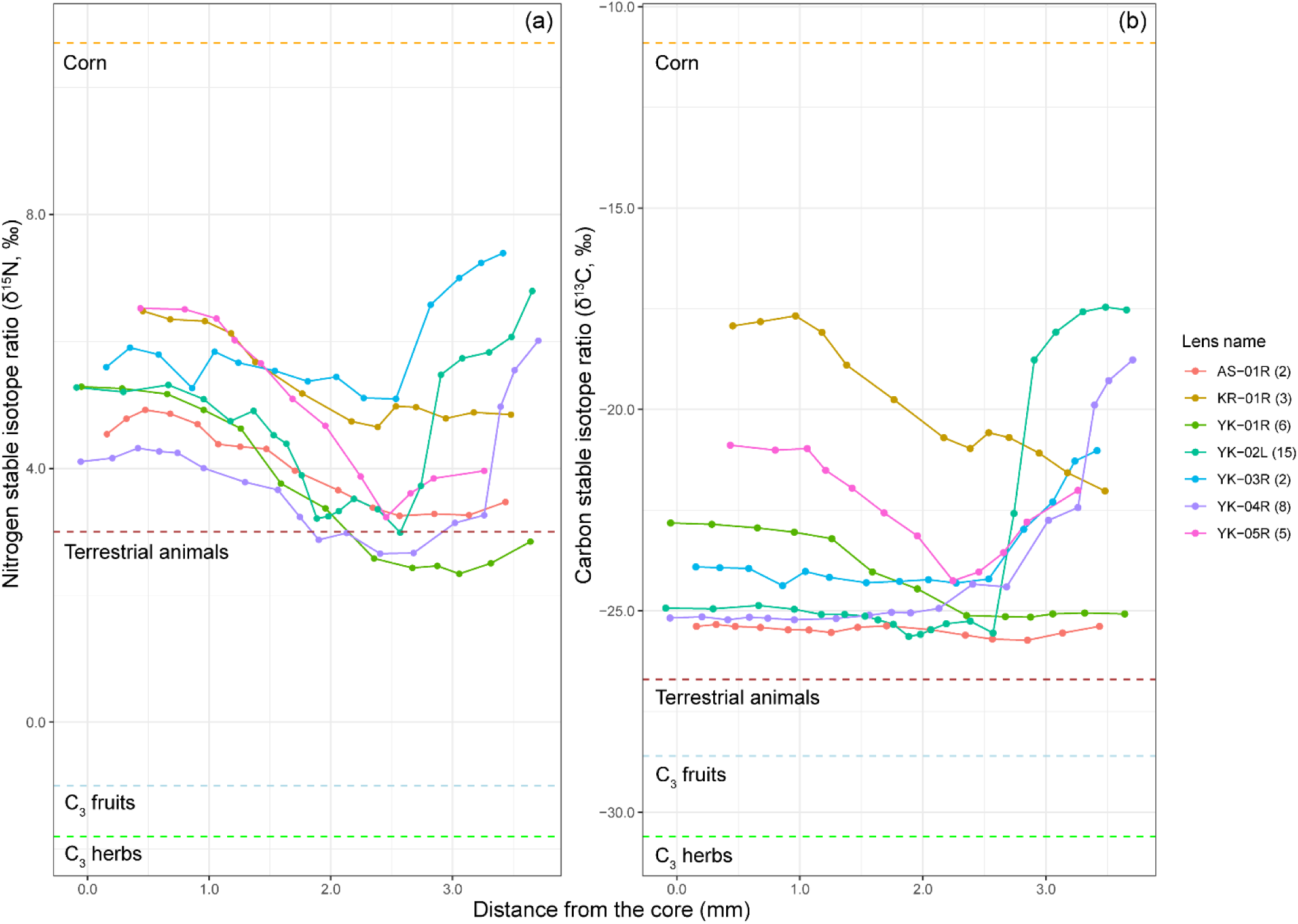
Fluctuation patterns of (a) nitrogen and (b) carbon stable isotope ratios (δ^15^N and δ^13^C) from eye lenses of seven brown bears. The solid lines represent the isotopic shifts within an individual lens. Dashed lines indicate the average δ^15^N and δ^13^C values of potential dietary sources of brown bears in the study region (C_3_ fruits, C_3_ herbs, terrestrial animals, and corn), as referenced by Matsubayashi et al. (2015) and Sakiyama et al. (2021). The numbers in parentheses in the legend indicate the age of each bear. Values less than zero on the x-axis indicate that the shaved segregated thickness slightly exceeds the lens radius.

For the four individuals from Yakumo (specifically YK-02L, YK-03R, YK-04R, and YK-05R), δ^15^N substantially increased from subsamples located >2.50 mm from the lens surface, approaching that of corn (Fig. 2a). The mean δ^15^N value of the outermost subsamples of these four individuals was 6.0‰ (± 1.5). For AS-01R, KR-01R, and YK-01R, no significant variation in δ^15^N was observed in the subsamples outside 2.50 mm from the core.

The mean δ^13^C of the most central subsample in each lens was higher for KR-01R and YK-05R (-19.4‰ ± 2.1) than for the other five individuals (-24.4‰ ± 1.1) (Fig. 2b). The δ^13^C of KR-01R decreased throughout their life, and that of YK-05R also decreased from the core to the subsample around 2.25 mm from the core. For YK-02L, YK-03R, YK-04R, and YK-05R, the δ^13^C largely increased and approached the corn value toward the outermost subsamples located >2.50 mm from the core (Fig. 2b). The δ^13^C increased by an average of -19.8‰ (± 2.0) in the outermost subsamples of the four individuals. The increasing δ^13^C trend of the four individuals was similar to the δ^15^N trend of these individuals (Fig. 2a). The δ^13^C of AS-01R and YK-01R were almost flat or decreased slightly before becoming constant from the core to the outer subsamples, and approached the values of terrestrial animals and C_3_ plants (Fig. 2b).

The δ^15^N and δ^13^C fluctuations of AS-01 and KR-01 were similar between the right and left eye lenses (Fig. 3), although the number of analysed subsamples and the distance from the core to each subsample were different. However, the left-right difference from around 1.00 mm to 2.25 mm from the core was relatively larger than the other subsamples for both pairs of lenses.

**Fig. 3.**
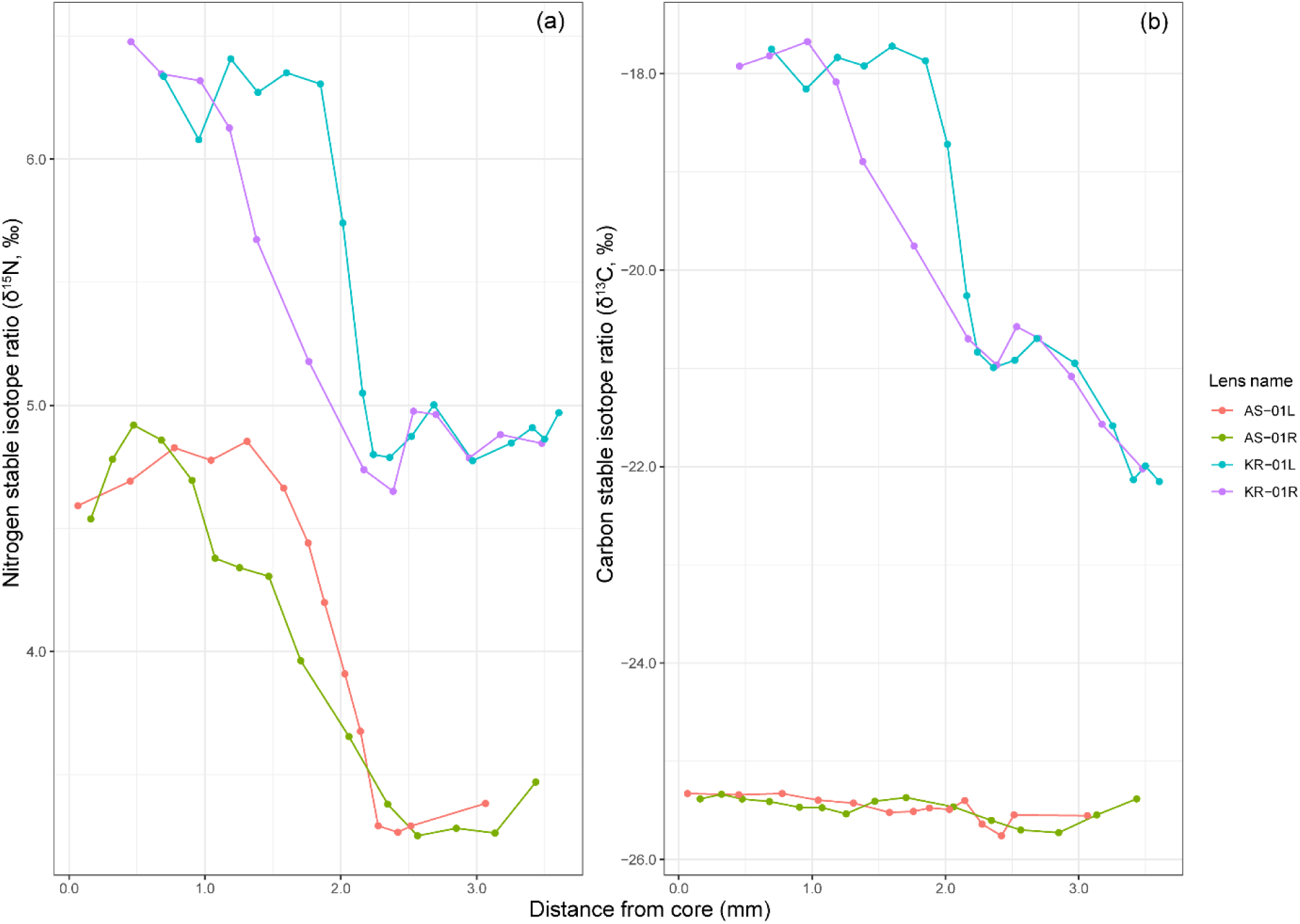
Fluctuation patterns of (a) nitrogen and (b) carbon stable isotope ratios (δ^15^N and δ^13^C) in two pairs of eye lenses. The solid lines represent the isotopic shifts in a single lens.

## 4. Discussion

The decreasing trend in δ^15^N in the sections from the inner half ( <2.50 mm from the core) of the eye lenses of all individuals may reflect changes in isotope ratios during lactation and weaning. Brown bear cubs rely solely on their mothers’ milk at birth (Jenkins et al., 2001). They then spend 1-2 years with their mothers (Shimozuru et al., 2017) and gradually transition to solid foods (Mazur & Seher, 2008). This dietary shift has been linked to changes in δ^15^N in other body tissues of young bears. For instance, grizzly bear (*Ursus arctos horribilis*) cubs demonstrated decreasing plasma δ^15^N levels before and after emerging from the den in their first three months (Jenkins et al. 2001). Similarly, polar bear (*Ursus maritimus*) cubs initially have higher plasma δ^15^N levels than their mothers immediately after leaving the den in the spring, but this difference diminishes by the summer (Polischuk et al., 2001). These findings support our hypothesis that retrospective isotope analysis using eye lenses can successfully reconstruct time-series isotopic histories from the lactation period.

The patterns of δ^13^C variation in all eye lens sections were largely classified into two categories. The first pattern showed δ^13^C stability in the inner half (<2.50 mm from the core) of the eye lenses, followed by a marked increase towards the outer sections (IDs: YK-02L, -03R, -04R). These individuals also displayed a concurrent increase in δ^15^N. Dent corn is the only diet component in the region that exhibits such elevated δ^13^C and δ^15^N (Matsubayashi et al., 2015; Sakiyama et al., 2021). Notably, three bears displaying this pattern originated from Yakumo, where dent corn is readily available. This isotopic trend confirms that sequential eye lens analysis accurately reconstructed individual diet history over several years. Until the end of weaning, these bears likely relied on non-corn sources, such as terrestrial animals and C_3_ plants, and shifted their diet towards dent corn in the post-weaning stage, as evidenced by the observed isotopic shift. The second major pattern involved consistently low δ^13^C (below -22.8‰) throughout all eye lens sections (YK-01R and AS-01R). These individuals also exhibited an initial decline in δ^15^N, which then plateaued. This pattern suggests a continuous dependence on non-corn diets throughout their lifetime.

Two individuals deviated from the above-mentioned patterns, exhibiting elevated δ^13^C and δ^15^N in their inner subsamples (<2.50 mm from the core). One individual (KR-01R) displayed consistently decreasing isotope ratios throughout its lifespan, while the other (YK-05R) exhibited increasing ratios in the further lens sections. Notably, the highest δ^13^C levels were detected near the innermost lens section for both individuals, indicating that the increase in δ^13^C likely originated from their mothers’ consumption of substantial quantities of corn rather than direct corn consumption by the bears themselves. In particular, since these individuals exhibited high levels of δ^13^C during the early life stages (at lens diameters <1.00 mm), it is inferred that this reflects the impact of a significant amount of corn consumption by the maternal bear in the year prior to giving birth. Additionally, the consistent decreasing trend in δ^13^C for KR-01R suggests that the bear was not dependent on a corn-heavy diet for most of its lifespan. Conversely, YK-05R’s data suggests that the bear did not rely on corn during lactation, but transitioned to a corn-based diet shortly after weaning. These contrasting isotopic patterns within brown bear lenses underscore the long-term variability in individual feeding behaviours, and demonstrate the efficacy of our novel method in identifying dietary differences at the individual level.

Another important finding of our study is the reconstruction of similar isotopic fluctuations from two eye lenses from the same individual. Similar results have also been shown in the eye lenses of other species, such as fish (Wallace et al., 2014). Our findings can increase sample availability when either eye lens is damaged during the killing or lens handling processes. Although our novel method was able to reconstruct similar isotopic trends from the right and left eye lenses of the same individual, the right-left isotopic differences were relatively large for subsamples located 1.00 to 2.25 mm from the core (Fig. 3). These differences could be attributed to the variability in the lens segregation process. In this study, a firmer texture was observed in the central region of the dried lens tissue than in the outer tissues. The nature of the sample might have contributed to the greater variability among subsamples. This aspect underscores the need for careful handling and consistent methodology when preparing lens samples for isotope analysis. Specifically, when segregating lens tissues, it is essential to standardise the tools and techniques used to ensure that sampling is conducted under uniform conditions.

Our data also provides insights into the temporal resolution of retrospective isotope analysis using mammalian eye lenses. For individuals that did not consume corn, as indicated by the time-series of isotope ratios (i.e. AS-01 and YK-01), their δ^15^N were stable around 2.50 mm from the core. This implies that subsamples obtained 2.50 mm from the lens core coincide with the transition from lactation to post-weaning, and that up to two years of dietary history was recorded from the core to these subsamples; in contrast, tissues obtained further than 2.50 mm from the lens core may contain information about the post-weaning diet of bears. In addition, in bears that consumed corn (i.e. YK-02 to -05), we did not detect any clear seasonal patterns in the lens isotope ratios, even though corn is a seasonal food source for Hokkaido brown bears (Hata et al., 2017). Thus, this method may not capture fine-scale dietary changes at the seasonal level after weaning. Furthermore, our analysis included brown bears aged 2 to 15 years, and we identified a weak correlation between eye lens diameter and age, suggesting that lens growth stabilises early in life. Our method mainly reflects time-series isotopic histories in the early life stages of bears. Future research could improve our methodology and increase its utility by correlating lens size with age.

In conclusion, this study proposes a novel and simple approach for retrospective isotope analyses of wild mammals using eye lenses, successfully reconstructing feeding behavioural histories from seven brown bears. Our findings enable the comparison of individual variations in the time-series behavioural pattern of mammals. These insights have significant implications for mammalian ecology and evolution as well as for conservation and management practices. Future research should aim to apply this method to a wider range of mammalian species, geographical regions, and other major stable isotope ratios, such as the sulphur stable isotope ratio (Crawford et al., 2008), thereby enhancing its utility. However, the limitations of this method, including the need for additional data on the lens growth patterns across different species, should be addressed by future studies. By overcoming these challenges, we can further improve the utility of retrospective isotope analysis in understanding individual-level behavioural variations in mammals.

## Acknowledgements

The authors are grateful to T. Sato and S. Ito for their assistance with the laboratory work. The authors would like to thank Y. Hangai provided mammalian eye lenses us to test the dissection and analysis, H. Mikami and H. Suzuki, who lent us the measurement devices, and S. Suzuki of the Yakumo Town Office provided important information regarding corn availability and bear-related crop damage in Yakumo Town. We would also like to thank all the people and organisations who assisted in the bear samples and data collection killed by nuisance control and hunting. This study was conducted with the support of the Joint Research Grant for the Environmental Isotope Study of Research Institute for Humanity and Nature.

## Author contribution statements

I. Tayasu, J. Matsubayashi and K. Miura conceived the ideas and designed methodology; H. Takafumi, H. Tsuruga, K. Miura, T. Mano, and Y. Shirane collected the samples and data; C. Yoshimizu, J. Matsubayashi, and K. Miura conducted isotope analysis; H. Takafumi and K. Miura analysed and illustrated the data; H. Tsuruga supervised the study. J. Matsubayashi and K. Miura led the writing of the original Draft; C. Yoshimizu, H. Takafumi, H. Tsuruga, I. Tayasu, J. Matsubayashi, K. Miura, T. Mano, and Y. Shirane conducted the review & editing of the draft. All authors contributed critically to the drafts and gave their final approval for publication.

## Data availability statement

We agree to archive the data associated with this manuscript should the manuscript be accepted to the “figshare (https://figshare.com/).”

