## Supplemental Information for "A novel method for fine-scale retrospective isotope analysis in mammals using eye lenses"

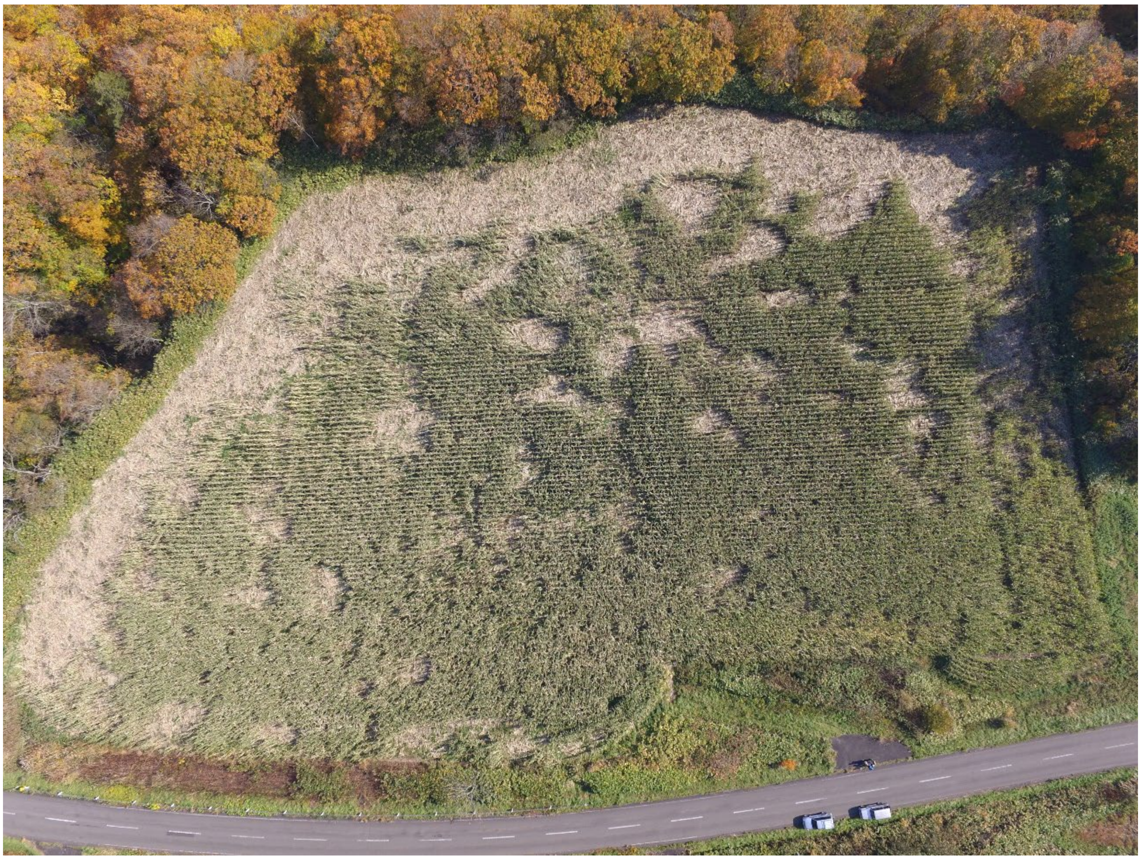

Fig. S1 Image of dent cornfields damaged by brown bears in Yakumo Town. White areas in the cornfield indicate bear-related damaged areas (Photo: Research Institute of Energy, Environment and Geology).

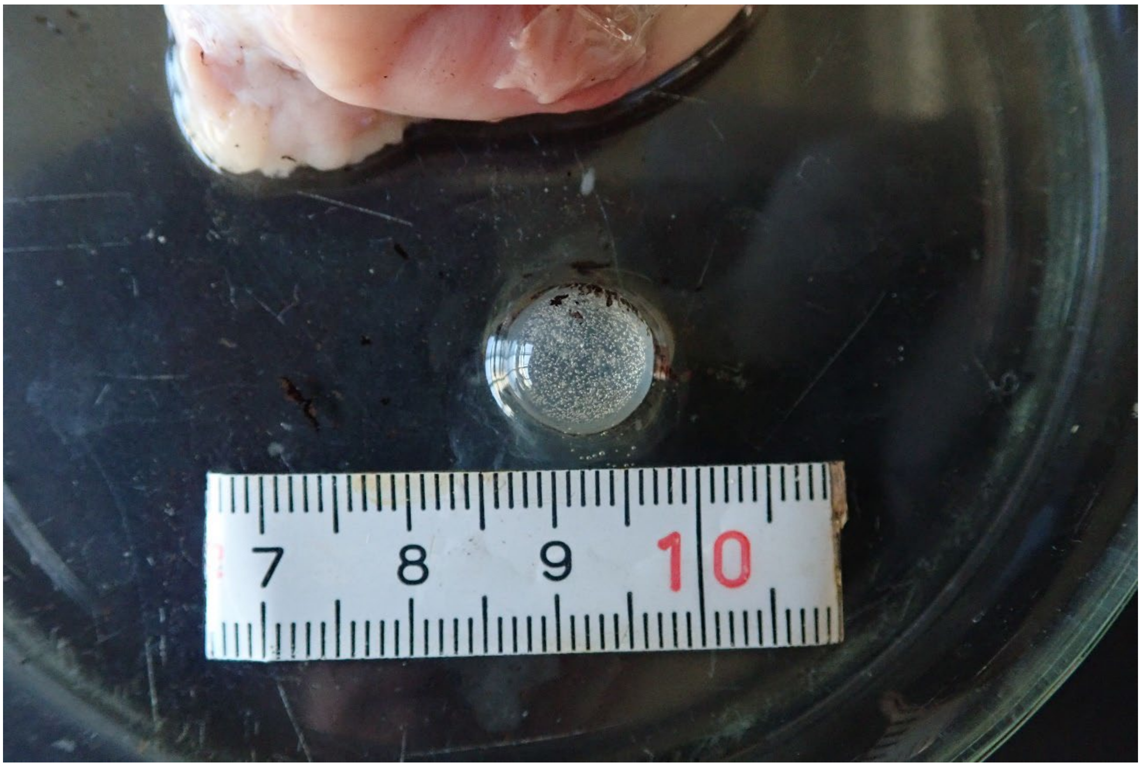


Fig. S2 Image of eye lens of brown bear in the raw state. The vitreous humor and lens capsule covering the eye lens have remained (Photo: Research Institute of Energy, Environment and Geology).
